## Supplementary Figures for "The multiplicity of Thioredoxin systems meets the specific needs of Clostridia"

**Table S1 List of strains used in this study**

| Strain | Genotype | Origin |
|---|---|---|
| ***E. coli*** | | |
| NEB10 | *Δ(ara-leu) 7697 araD139 fhuA ΔlacX74 galK16 galE15 e14-* ϕ*80*d*lacZΔM15 recA1 relA1 endA1 nupG rpsL* ($Str^R$) *rph spoT1 Δ(mrr-hsdRMS-mcrBC)* | BioLabs |
| HB101(RP4) | *supE*44 *aa*14 *galK*2 *lacY*1 Δ(*gpt-proA*) 62 *rpsL*20 ($Str^R$)*xyl-5 mtl-1 recA*13 Δ(*mcrC-mrr*) *hsdS*B($r_B^-m_B^-$) RP4 ($Tra^+$ IncP $Ap^R$ $Km^R$ $Tc^R$) | Laboratory stock |
| ***C. difficile*** | | |
| 630Δ*erm* | | Laboratory stock |
| E1 | | Laboratory stock |
| CDIP229 | 630Δ*erm sigB*::*erm* | Kint *et al.*, 2017 |
| CDIP529 | 630Δ*erm trxB1*::*erm* | This work |
| CDIP1456 | 630Δ*erm* pMTL84121 | Laboratory stock |
| CDIP1461 | 630Δ*erm* Δ*trxB2* | This work |
| CDIP1486 | 630Δ*erm trxB1*::*erm* Δ*trxB2* | This work |
| CDIP1632 | 630Δ*erm trxB1*::*erm* Δ*trxB2* pMTL84121 | This work |
| CDIP1636 | 630Δ*erm trxB1*::*erm* Δ*trxB2* pMTL84121-P-*trxB2* | This work |
| CDIP1640 | 630Δ*erm trxB1*::*erm* Δ*trxB2* pMTL84121-P-*trxB1* | This work |
| CDIP1796 | 630Δ*erm* Δ*trxA1B1* | This work |
| CDIP1797 | 630Δ*erm* Δ*trxA2* | This work |
| CDIP1811 | 630Δ*erm* Δ*trxA1* | This work |
| CDIP1812 | 630Δ*erm* Δ*trxA1* Δ*trxA2* | This work |
| CDIP1814 | 630Δ*erm* Δ*trxA1* Δ*trxA2* pMTL84121-P-*trxA1* | This work |
| CDIP1815 | 630Δ*erm* Δ*trxA1* Δ*trxA2* pMTL84121-P-*trxA2* | This work |
| CDIP1823 | 630Δ*erm* Δ*trxA1* Δ*trxA2* pMTL84121 | This work |
| CDIP1884 | 630Δ*erm trxB1*::*erm* Δ*trxB2* pMTL84121-P-*trxB4* (E1 strain) | This work |
| CDIP1924 | 630Δ*erm* Δ*trxA3* | This work |
| CDIP1925 | 630Δ*erm* Δ*trxA1* Δ*trxA3* | This work |
| CDIP1933 | 630Δ*erm* Δ*trxA2* Δ*trxA3* | This work |
| CDIP1939 | 630Δ*erm* Δ*trxA1* Δ*trxA2* Δ*trxA3* | This work |
| CDIP1967 | 630Δ*erm* Δ*trxA1* Δ*trxA2* Δ*trxA3* pMTL84121 | This work |
| CDIP1968 | 630Δ*erm* Δ*trxA1* Δ*trxA2* Δ*trxA3* pMTL84121-P-*trxA1* | This work |
| CDIP1969 | 630Δ*erm* Δ*trxA1* Δ*trxA2* Δ*trxA3* pMTL84121-P-*trxA3* | This work |
| CDIP1970 | 630Δ*erm* Δ*trxA1* Δ*trxA2* Δ*trxA3* pMTL84121-P-*trxA2* | This work |
| CDIP2113 | 630Δ*erm* Δ*grdAB* | This work |
| CDIP1982 | 630Δ*erm* pFT47-$P_{trxA1B1}$-$FAST^{CD}$ | This work |
| CDIP1983 | 630Δ*erm sigB*::*erm* pFT47-$P_{trxA1B1}$-$FAST^{CD}$ | This work |
| CDIP2157 | 630Δ*erm sigB*::*erm* pFT47-$FAST^{CD}$ | This work |
| CDIP2075 | 630Δ*erm* pFT47-$P_{trxA1B1}$-TrxA1'-$FAST^{CD}$ | This work |
| CDIP2203 | 630Δ*erm sigF*::*erm* pFT47-$P_{trxA1B1}$-$FAST^{CD}$ | This work |
| CDIP2205 | 630Δ*erm sigG*::*erm* pFT47-$P_{trxA1B1}$-$FAST^{CD}$ | This work |

P: promoter, *erm*: erythromycin resistance gene. *trxA1 = CD1690*, *trxA2 = CD3033*, *trxA3 = CD2355, trxB1 = CD1691*, *trxB2 = CD2117*

**Table S2 List of plasmids**

| Plasmid | Characteristics | Origin |
|---|---|---|
| pMTL007 | Plasmid for gene inactivation through Clostron | Laboratory stock |
| pMSR | Plasmid for gene deletion through allelic chromosomic exchange (ACE) | Laboratory stock |
| pMTL84121 | Replicative plasmid in *E. coli* and *C. difficile* and conjugation-transmissible | Laboratory stock |
| pGEMTeasy | Plasmid for TA cloning for sequencing for 5'RACE | Promega |
| pDIA6190 | pMTL007-Cdi-*trxB1-36a* | This work |
| pDIA6919 | pMSR-ACE Δ*trxB2* | This work |
| pDIA7025 | pMTL84121-P-*trxA1-trxB1* | This work |
| pDIA7042 | pMTL84121-P-*trxB1* | This work |
| pDIA7050 | pMTL84121-P-*trxB2* | This work |
| pDIA7108 | pMSR-ACE Δ*trxA1-trxB1* | This work |
| pDIA7112 | pMSR-ACE Δ*trxA2* | This work |
| pDIA7113 | pMTL84121-P-*trxA2-trxB4* (strain E1) | This work |
| pDIA7118 | pMSR-ACE Δ*trxA1* | This work |
| pDIA7122 | pMTL84121-P-*trxA2* | This work |
| pDIA7129 | pMTL84121-P-*trxA1* | This work |
| pDIA7156 | pMTL74121-P-*trxB4* (strain E1) | This work |
| pDIA7162 | pMTL84121-P-*grdX-trxB3-trxA3* | This work |
| pDIA7163 | pMSR-ACE Δ*trxA3* | This work |
| pDIA7164 | pMTL84121-P-*trxA3* | This work |
| pDIA7190 | pFT47-$FAST^{CD}$ | This work |
| pDIA7194 | pFT47-$P_{trxA1B1}$-$FAST^{CD}$ | This work |
| pDIA7236 | pFT47-$P_{trxA1B1}$-*trxA1'*-$FAST^{CD}$ | This work |
| pDIA7281 | pMSR-ACE Δ*grdAB* | This work |

ACE: allelic chromosomic exchange, P: promoter.

**Table S3 List of oligonucleotides**

| Primer | Sequence | Characteristics |
|---|---|---|
| EBSu | CGAAATTAGAAACTTGCGTTCAGTAAAC | Clostron *trxB1* |
| IMV690 | AAAAAAGCTTATAATTATCCTTAGCTGGCCCACTGGTGCGCCCAGATAGGGTG | |
| IMV691 | CAGATTGTACAAATGTGGTGATAACAGATAAGTCCCACTGCCTAACTTACCTTTCTTTGT | |
| IMV692 | TGAACGCAAGTTTCTAATTTCGATTCCAGCTCGATAGAGGAAAGTGTCT | |
| IMV1113 | TGGTCATGAGATTATCAAAAGGTTGGAACATAAAGACCACCATCATT | ACE mutagenesis *trxB2* |
| IMV1114 | TCCAGCACCAATTACAATGATATCT | |
| IMV1115 | AGATATCATTGTAATTGGTGCTGGAGGAGCTATTGCAGCCGTTCAA | |
| IMV1116 | ATCGTAGAAATACGGTGTTTTTGGAGAAATAGTTGTTATAGCTGGTA | |
| CA5 | TGGTCATGAGATTATCAAAAGGGATGAGTTACTTCCAGATAGAAG | ACE mutagenesis *trxA1* |
| CA6 | TTTTAAATCCTCCTATTAAACTTTATTATC | |
| CA7 | GATAATAAAGTTTAATAGGAGGATTTAAAAGTATCTACAGTACCAACTATG | |
| CA19 | ATCGTAGAAATACGGTGTTTTTGTACTCTTAATTAGATTTTTCAACC | |
| CA9 | TGGTCATGAGATTATCAAAAGGGCAGCAGGTCTTTATG | ACE mutagenesis *trxA2* |
| CA10 | TTTCTTATTCCCCCTTGAAA | |
| CA11 | TTTCAAGGGGGAATAAGAAAGGTCTTCCAACTATGGCT | |
| CA12 | ATCGTAGAAATACGGTGTTTTTGAAACACAGTTACCACTTAC | |
| CA1 | TGGTCATGAGATTATCAAAAGGGTATTGAACTCCAATATATAAAATG | ACE mutagenesis *trxA3* |
| CA2 | AATATACATCTCCTTTATTATCTTC | |
| CA3 | GAAGATAATAAAGGAGATGTATATTGGAAAACCTGTAGATAGATTAAT | |
| CA4 | ATCGTAGAAATACGGTGTTTTTGTAATGTATCTCTCGTTACAG | |
| CA13 | GATAATAAAGTTTAATAGGAGGATTTAAAAGCTGGTCAAGGG | ACE *trxA1-trxB1* |
| CA14 | ATCGTAGAAATACGGTGTTTTTGTAAACTTTTATTCCCATTCC | |
| CM13 | GATCGGTCTTGCCTTGCTC | Amplification of pMTL84121 for cloning through Gibson Assembly |
| IMV993 | CTGGCGTTACCCAACTTAATCG | |
| IMV1183 | CCGCTCGAGATATTGGAACTACATTTGAATTG | Amplification of P-*trxA1-trxB1* |
| IMV1151 | GGGGATCCTTAAAAATAAAAAACTGTCTTGA | |
| IMV1200 | AAAATGGGTGAGTATTATGAGA | Inverse PCR to obtain P-*trxB1* |
| IMV1201 | CAGATGTATTTATAACTTTTGC | |
| IMV1331 | GGACCACTGCCGATTATAGCT | Inverse PCR to obtain P-*trxA1* |
| CA24 | GAGCAAGGCAAGACCGATCGGGAAAAAATGTCAACTATATTAAG | Amplification of P-*trxA2* |
| CA25 | CGATTAAGTTGGGTAACGCCAGCATTAGCAATTAGACACTCATTTG | |
| IMV1202 | GAGCAAGGCAAGACCGATCACAGCAGCATTAGTTTTGGTTAT | Amplification of P-*trxB2* |
| IMV1203 | CGATTAAGTTGGGTAACGCCAGTGAACATGATAGATTAAGATATAAC | |
| IMV1235 | GAGCAAGGCAAGACCGATCGAAAATACAAGAAATATGCAATATCAA | Amplification of P-*grdX-trxB3-trxA3* |
| CA43 | CGATTAAGTTGGGTAACGCCAGCATAAAAACCTTTTTCTTCTTATCTTC | |
| IMV1300 | TCTGCCTTGAATTCTTCTTTTACT | Inverse PCR to obtain P-*trxA3* |
| IMV1389 | GGGAACTAGCAATCTGTAAATTAAG | |
| CA26 | GAGCAAGGCAAGACCGATCGCTTATCATAATATTAAAGCCAG | Amplification of P-*trxA2-trxB4* (E1) |
| CA27 | CGATTAAGTTGGGTAACGCCAGGAATCCACAATCAATAACAAC | |
| CA75 | AATTTACATCTCCTTTATTATTTTC | Inverse PCR to obtain P-*trxB4* |
| CA76 | TTAAAATAAATCATATATAAACAAAAAAC | |
| AAP | GGCCACGCGTCGACTAGTACGGGIIGGGIIGGGIIG | Abridged Anchor Primer |
| IMV710 | TTATCATTACCAAACATGA | 5'RACE mRNA amplification *trxB1* |
| IMV1209 | GTTTCTACTGGTTTTCCATCTTT | 5'RACE TA Cloning *trxB1* |
| IMV1156 | CACCTGAAGAAGTTTTTACAAC | 5'RACE mRNA amplification *trxB2* |
| IMV1157 | TCAACACCTTGAGCAACTGC | 5'RACE TA Cloning *trxB2* |
| IMV567 | CTCCAAGTGCATTGGTTCCT | qPCR *trxB1* |
| IMV568 | AATCTCCAGCAGCAAAGCAT | |
| IMV571 | TCAAGCAGTTGCTCAAGGTG | qPCR *trxB2* |
| IMV572 | AATTCTTGTTCCCCTGCACA | |

| | | |
|---|---|---|
| IMV1239 | TGGTTGTCCTGGAGAAAAAGA | qPCR *trxB3* |
| IMV1240 | ATGCACTGTCTCCTCCACCT | |
| IMV1274 | TTTCTTTGCGACTTGGTGTG | qPCR *trxA3* |
| IMV1275 | GTCCACCTTCAAGAAGTTTGCT | |
| IMV1237 | TATGAGCGATGAAGGGCATT | qPCR *CD3605.1* |
| IMV1238 | AGCTGAAACTGGACATCCTTCT | |
| BD3 | TTTTGTTGTGTCTATGAACCTTTGT | qPCR *gyrA* |
| BD4 | TCCTTTACCAGCTCTTATTTGACTT | |
| BD9 | GTTAAATGGGATAGAAGAGGTTGCT | qPCR *ccpA* |
| BD10 | TATACCTTCCACTTGTTTTGCTCTC | |
| IMV1103 | CCGAGCTCGAATTCGTAATCATGGT | Amplification of pFT47 for cloning through Gibson Assembly |
| IMV1140 | CCGCTCGAGCATAAAATCATCCTCTCTTATATT | |
| IMV1105 | ACCATGATTACGAATTCGAGCTCGGACATTTGAATTGTCAATACCA | Amplification of $P_{trxA1B1}$ |
| IMV1443 | CATGGTATTTTCCTCCTTTCTCCAGATGTATTTATAACTTTTGC | |
| IMV1499 | AACCAAAGGCTACATGCTCCATTAAATATGATTTTACTTGCATTTCAATTTT | Amplification of $P_{trxA1B1}$-*trxA1’* |

**Fig. S1. Conservation of protein sequences of Trx partners and of genetic organization of the *grd* operon.** (A) Alignment of TrxB sequences from diverse bacteria. Alignment was performed using MAFFT software [110]. The CxxC active motif is indicated by a red box and the NAD(P)H binding motifs by a green box. (B) Distance tree of TrxBs was obtained using the neighbor-joining method. FFTRs cluster in orange and NTRs in green. The atypical TrxB from *Desulfovibrio vulgaris* [112] was used to root the tree. Bootstraps are indicated on the branches. (C) Alignment of TrxA sequences from diverse bacteria. Alignment was performed using MAFFT. CxxC active domain is indicated by a red box, the classical GP site is highlighted in green and the atypical V/EP site in blue. (D) Synteny of Clostridial *grd* operons. Sequence of *grd* operons from proteolytic Clostridia were analyzed using the MicroScope platform [57].

**Fig. S2. Promoters of the *trx* genes and regulation of the *trxB1* gene and of genes encoding its associated ferredoxin.** (A) Promoter identification through 5'RACE using RNA extracted from exponentially growing cells of strain 630Δ*erm*. The TSS (+1) is indicated in red. Upstream this TSS, $\sigma^B$ boxes are represented in orange. (B-C) Expression of the *trxB1* gene and the *CD3605.1* gene encoding a ferredoxin was monitored by qRT-PCR in (B) WT strain after 24 h of growth in TY medium in anaerobiosis or at 1% $O_2$ or in (C) WT strain and *sigB* mutant after 4.5 h of growth in TY. Experiments were performed in at least 6 biological replicates. Mean and SD are shown. One sample t-tests were used with comparison of the fold change to 1. *: p-value<0.05, ** <0.01, *** <0.001.

**Fig. S3. Growth curves and survival of *trx* mutants.** (A) Growth curves of the different *trx* mutants. Growth was monitored in a 96-well plate using an initial bacterial suspension at $OD_{600nm}$ 0.05 in TY for 24 h at 37°C. Experiments were performed in 5 biological replicates. Mean and SD are shown. The first panel presents all curves represented in other panels as individual curves. (B, C) Survival of (B) *trxA* and (C) *trxB* mutants. A bacterial suspension at $OD_{600nm}$ 0.05 in TY was prepared. Total bacteria were numerated daily over 2 days by plating serial dilutions on TY Tau plates. Experiments were performed in 5 biological replicates. Mean and SEM are shown. Multiple unpaired t-tests were performed. *: p-value<0.05, ** <0.01, *** <0.001.

**Fig. S4. Stress tolerance of complemented strains.** (A, B) Strains were serially diluted, plated in duplicate on TY Tau plates and incubated either in anaerobiosis or in hypoxia at (A) 1% $O_2$ or (B) 0.1% $O_2$ for 64 h. Survival was then normalized by doing the ratio of the mutant *vs* the WT. (C, D) Strains were serially diluted and plated on TY and on (C) TY + DEA NONOate 750 µM or (D) TY + HClO 0.1% and incubated for 24 h. Mean and SD are shown. Experiments were performed in 5 biological replicates. For all assays, ordinary one-way ANOVA were performed followed by Dunett's multiple comparison tests. *: p-value<0.05, ** <0.01, *** <0.001 and **** <0.0001.

**Fig. S5. Conservation of the *trxB4* gene and the promoter of the *trxA2-trxB4* operon.** (A) Comparison of the *C. difficile* core genome and *trxB4* evolution. The tanglegram representing the *trxB4* tree (right) and the core-genome tree pruned to contain only genomes present in the *trxB4* tree (left). The branch lengths are measured in substitutions *per* site and are on the same scale. The only difference between the two trees is that the *trxB4* one is less resolved, which is explained by the limited phylogenetic signal in only one gene used for its reconstruction. The figure is produced with Dendroscope [109]. (B) Alignment of *trxA2* promoter region in the 630Δ*erm* and the E1 strains. Regions corresponding to the 150 bp upstream the *trxA2* start codon (ATG) and the following 100 bp from the 630Δ*erm* and the E1 strains were aligned using the MAFFT software [110]. The ATG, the TSS and the $\sigma^A$ promoter [46] are indicated. Jalview software [111] was used for visualization.

**Fig. S6. Bile acids tolerance of complemented strains.** Strains were serially diluted and plated on TY and on (A) TY + DOC 0.03%, (B) TY + CHO 0.4%, (C) TY + Triton X-100 0.01%, (D) TY + SDS 0.003% or (E) TY + diamide 0.02% and incubated for 24 h. Survival was calculated by doing the ratio between CFUs in the last dilution with stress and CFUs in the last dilution without stress. Mean and SD are shown. Experiments were performed in 5 biological replicates. For all assays, ordinary one-way ANOVA were performed followed by Dunett's multiple comparison test. *: p-value<0.05, ** <0.01, *** <0.001 and **** <0.0001.

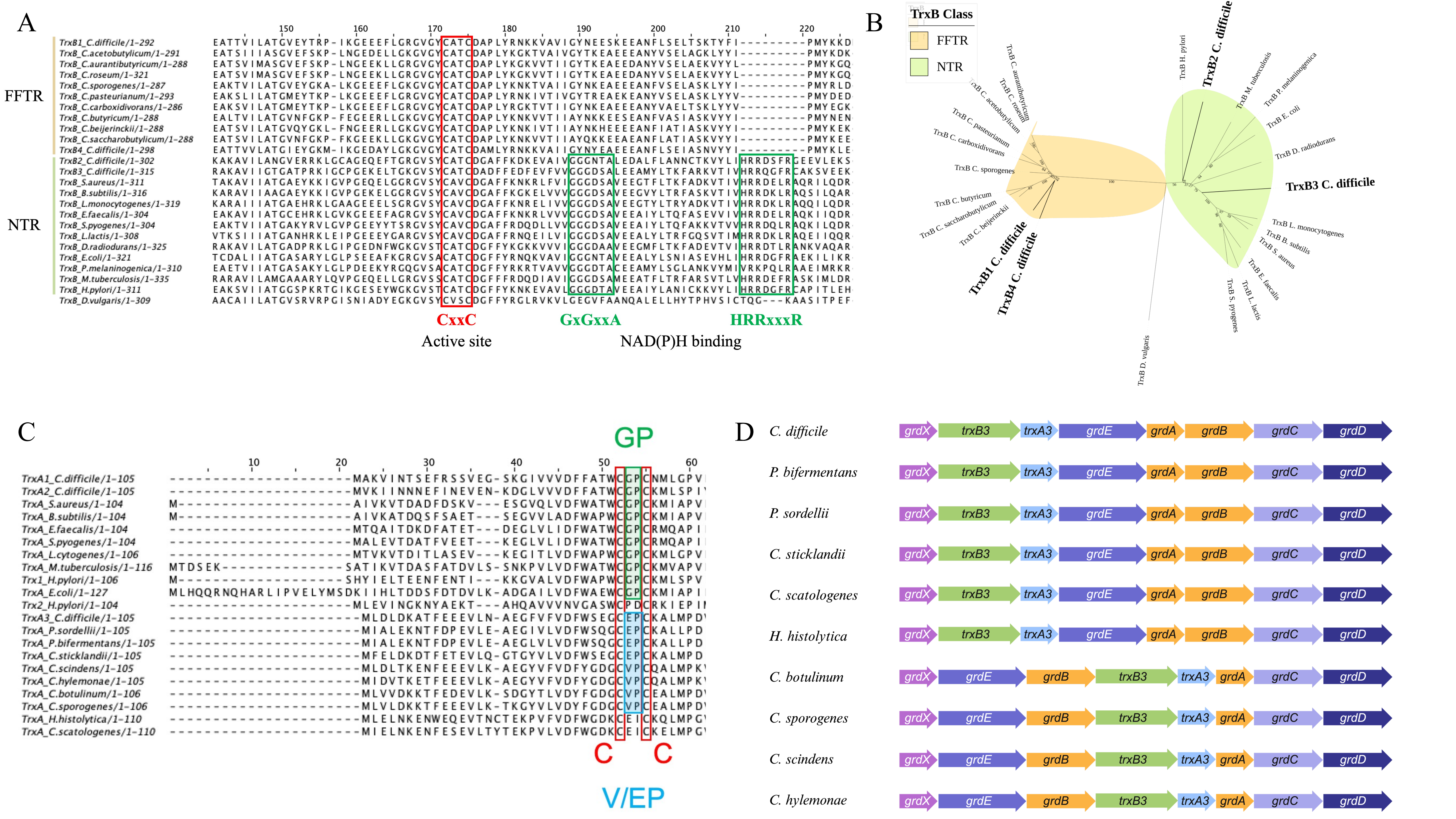
A
FFTR
NTR
TrxB1_C.difficile/1-292
TrxB_C.acetobutylicum/1-291
TrxB_C.aurantibutyricum/1-288
TrxB_C.roseum/1-321
TrxB_C.sporogenes/1-287
TrxB_C.pasteurianum/1-293
TrxB_C.carboxidivorans/1-286
TrxB_C.butyricum/1-288
TrxB_C.beijerinckii/1-288
TrxB_C.saccharobutylicum/1-288
TrxB4_C.difficile/1-298
TrxB2_C.difficile/1-302
TrxB3_C.difficile/1-315
TrxB_S.aureus/1-311
TrxB_B.subtilis/1-316
TrxB_L.monocytogenes/1-319
TrxB_E.faecalis/1-304
TrxB_S.pyogenes/1-304
TrxB_L.lactis/1-308
TrxB_D.radiodurans/1-325
TrxB_E.coli/1-321
TrxB_P.melaninogenica/1-310
TrxB_M.tuberculosis/1-335
TrxB_H.pylori/1-311
TrxB_D.vulgaris/1-309
CxxC
Active site
GxGxxA
HRRxxxR
NAD(P)H binding
B
TrxB Class
FFTR
NTR
TrxB1 C. difficile
TrxB4 C. difficile
TrxB2 C. difficile
TrxB3 C. difficile
TrxB D. vulgaris
C
GP
C
C
V/EP
TrxA1_C.difficile/1-105
TrxA2_C.difficile/1-105
TrxA_S.aureus/1-104
TrxA_B.subtilis/1-104
TrxA_E.faecalis/1-104
TrxA_S.pyogenes/1-104
TrxA_L.cytogenes/1-106
TrxA_M.tuberculosis/1-116
Trx1_H.pylori/1-106
TrxA_E.coli/1-127
Trx2_H.pylori/1-104
TrxA3_C.difficile/1-105
TrxA_P.sordellii/1-105
TrxA_P.bifermentans/1-105
TrxA_C.sticklandii/1-105
TrxA_C.scindens/1-105
TrxA_C.hylemonae/1-105
TrxA_C.botulinum/1-106
TrxA_C.sporogenes/1-106
TrxA_H.histolytica/1-110
TrxA_C.scatologenes/1-110
D
C. difficile
P. bifermentans
P. sordellii
C. sticklandii
C. scatologenes
H. histolytica
C. botulinum
C. sporogenes
C. scindens
C. hylemonae
grdX
trxB3
trxA3
grdE
grdA
grdB
grdC
grdD

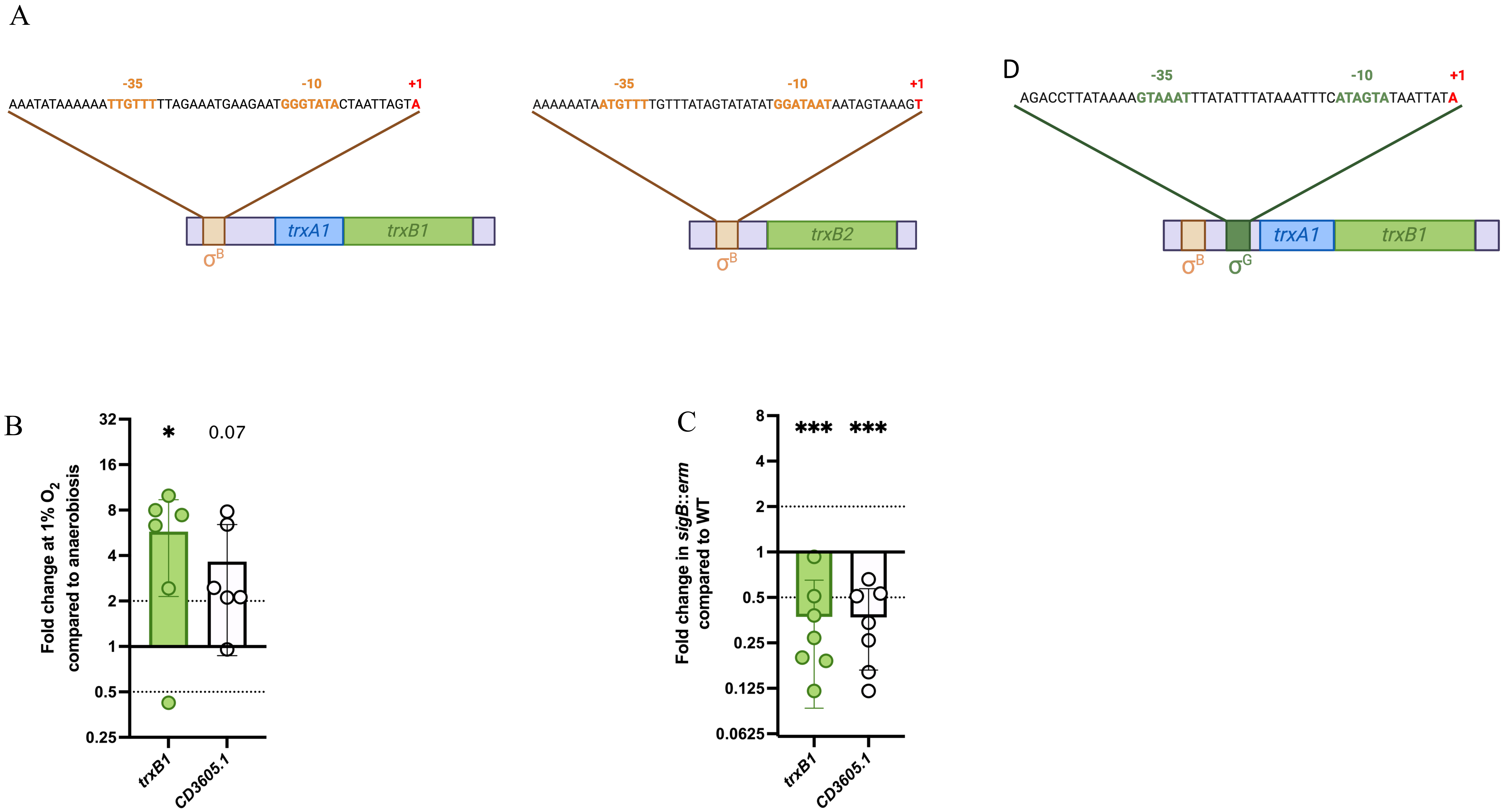
A
-35
-10
+1
AAATATAAAAAATTGTTTTTAGAAATGAAGAATGGGTATACTAATTAGTA
trxA1
trxB1
σB
-35
-10
+1
AAAAAAATAATGTTTTGTTTATAGTATATATGGATAATAATAGTAAAGT
trxB2
σB
D
-35
-10
+1
AGACCTTATAAAAGTAAATTTATATTTATAAATTTCATAGTATAATTATA
σB
σG
trxA1
trxB1
B
Fold change at 1% O2 compared to anaerobiosis
32
16
8
4
2
1
0.5
0.25
*
0.07
trxB1
CD3605.1
C
Fold change in sigB::erm compared to WT
8
4
2
1
0.5
0.25
0.125
0.0625
***
***
trxB1
CD3605.1

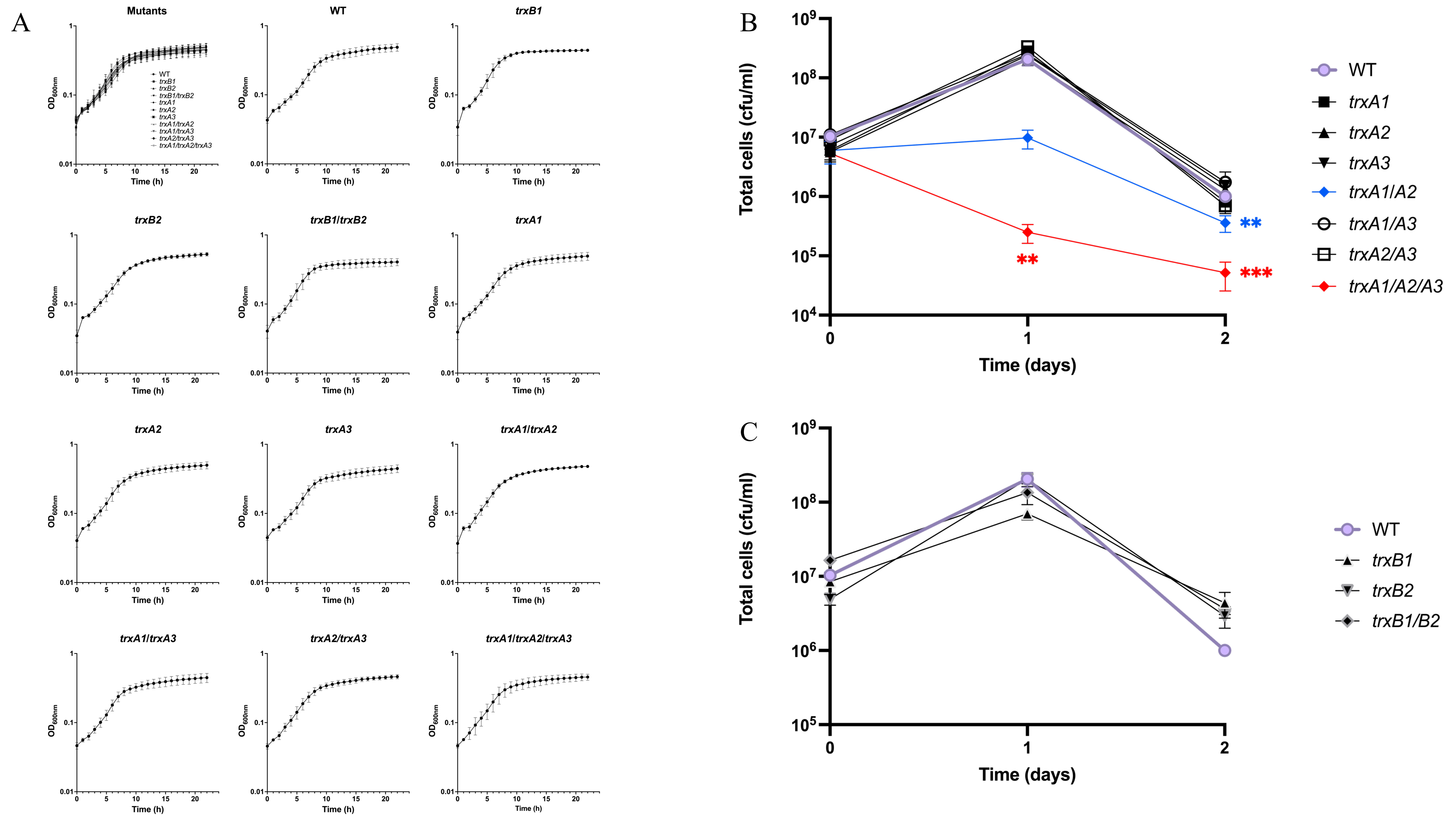
A
Mutants
WT
trxB1
trxB2
trxB1/trxB2
trxA1
trxA2
trxA3
trxA1/trxA2
trxA1/trxA3
trxA2/trxA3
trxA1/trxA2/trxA3
OD600nm
Time (h)
B
Total cells (cfu/ml)
Time (days)
WT
trxA1
trxA2
trxA3
trxA1/A2
trxA1/A3
trxA2/A3
trxA1/A2/A3
**
***
C
Total cells (cfu/ml)
Time (days)
WT
trxB1
trxB2
trxB1/B2

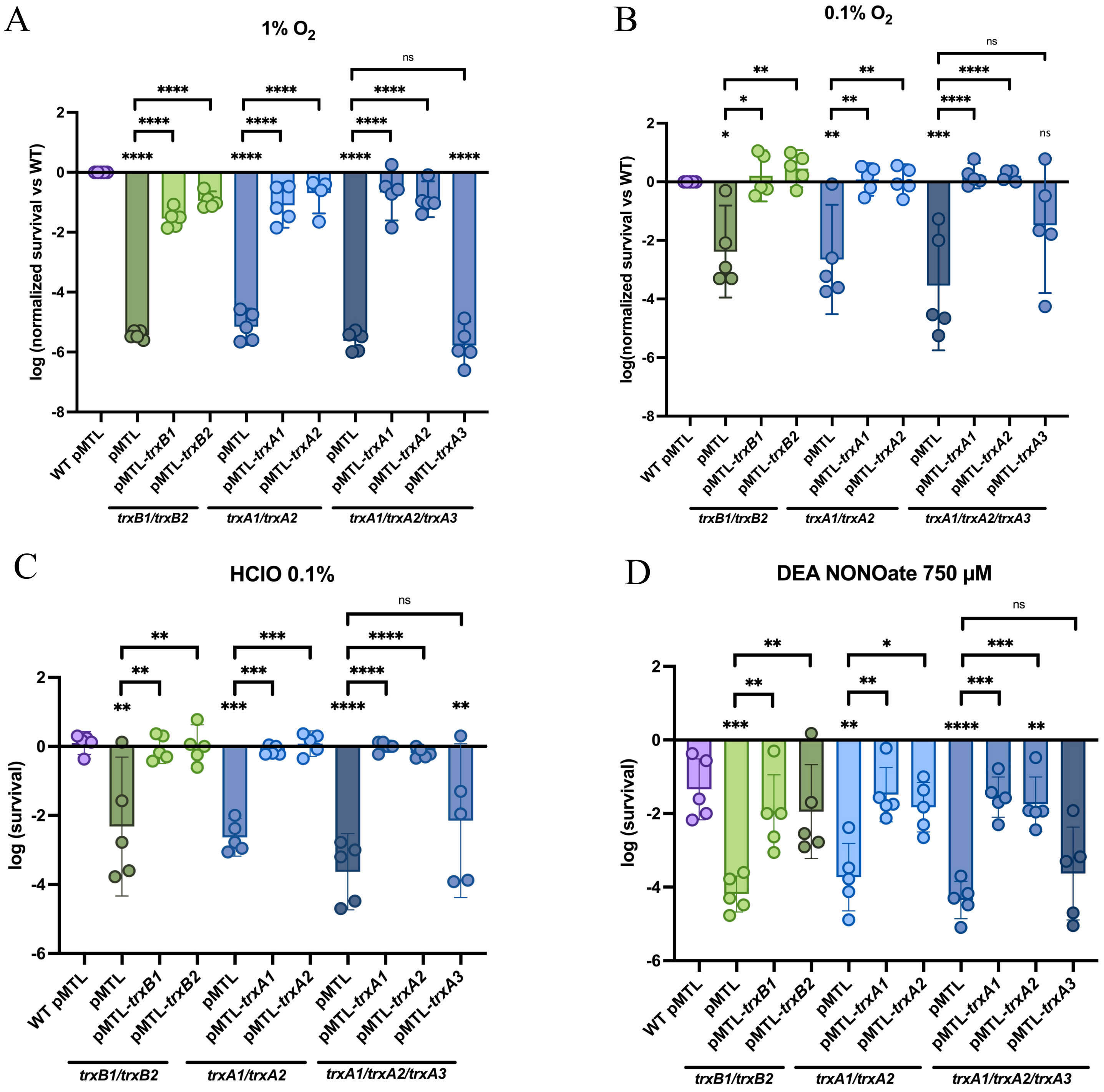
A
1% $O_2$
B
0.1% $O_2$
C
HClO 0.1%
D
DEA NONOate 750 µM
log (normalized survival vs WT)
log(normalized survival vs WT)
log (survival)
WT pMTL
pMTL
pMTL-*trxB1*
pMTL-*trxB2*
pMTL-*trxA1*
pMTL-*trxA2*
pMTL-*trxA3*
*trxB1/trxB2*
*trxA1/trxA2*
*trxA1/trxA2/trxA3*
ns

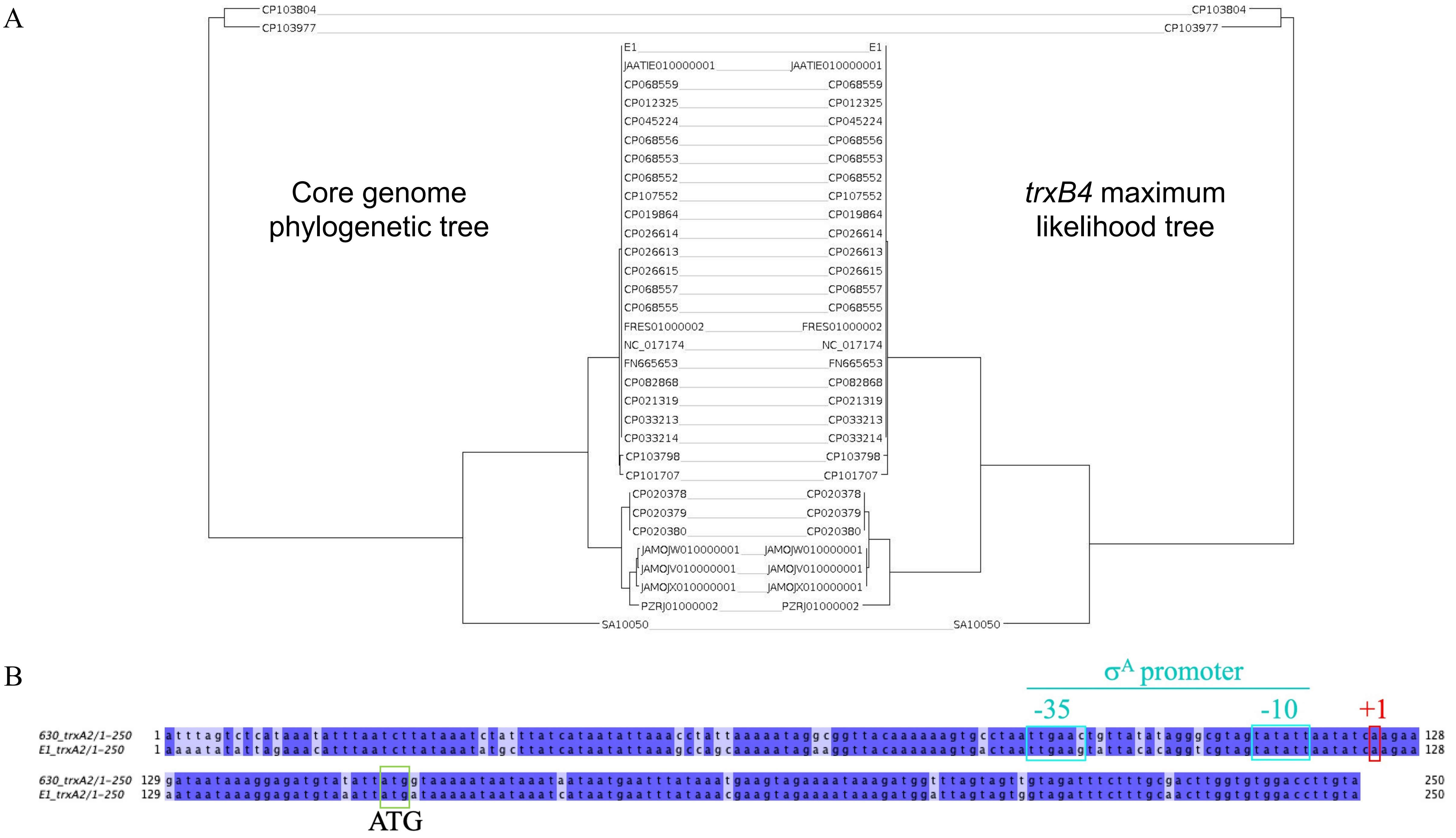

A
CP103804
CP103977
E1
JAATIE010000001
CP068559
CP012325
CP045224
CP068556
CP068553
CP068552
CP107552
CP019864
CP026614
CP026613
CP026615
CP068557
CP068555
FRES01000002
NC_017174
FN665653
CP082868
CP021319
CP033213
CP033214
CP103798
CP101707
CP020378
CP020379
CP020380
JAMOJW010000001
JAMOJV010000001
JAMOJX010000001
PZRJ01000002
SA10050
Core genome phylogenetic tree
trxB4 maximum likelihood tree
B
σA promoter
-35
-10
+1
630_trxA2/1-250 1 atttagtctcataaatatttaatcttataaatctatttatcataatattaaacctattaaaaataggcggttacaaaaaagtgcctaattgaactgttatatagggcgtagtatattaatatcaagaa 128
E1_trxA2/1-250 1 aaaatatattagaaacatttaatcttataaatatgcttatcataatattaaagccagcaaaaatagaaggttacaaaaaagtgactaattgaaggtattacacaggtcgtagtatattaatatcaagaa 128
630_trxA2/1-250 129 gataataaaggagatgtatattatggtaaaaataataaataataatgaatttataaatgaagtagaaaataaagatggttagtagttgtagatttctttgcgacttggtgtggaccttgta 250
E1_trxA2/1-250 129 aataataaaggagatgtaaattatgataaaaataataaatcataatgaatttataaacgaagtagaaaataaagatggattagtagtggtagatttctttgcaacttggtgtggaccttgta 250
ATG

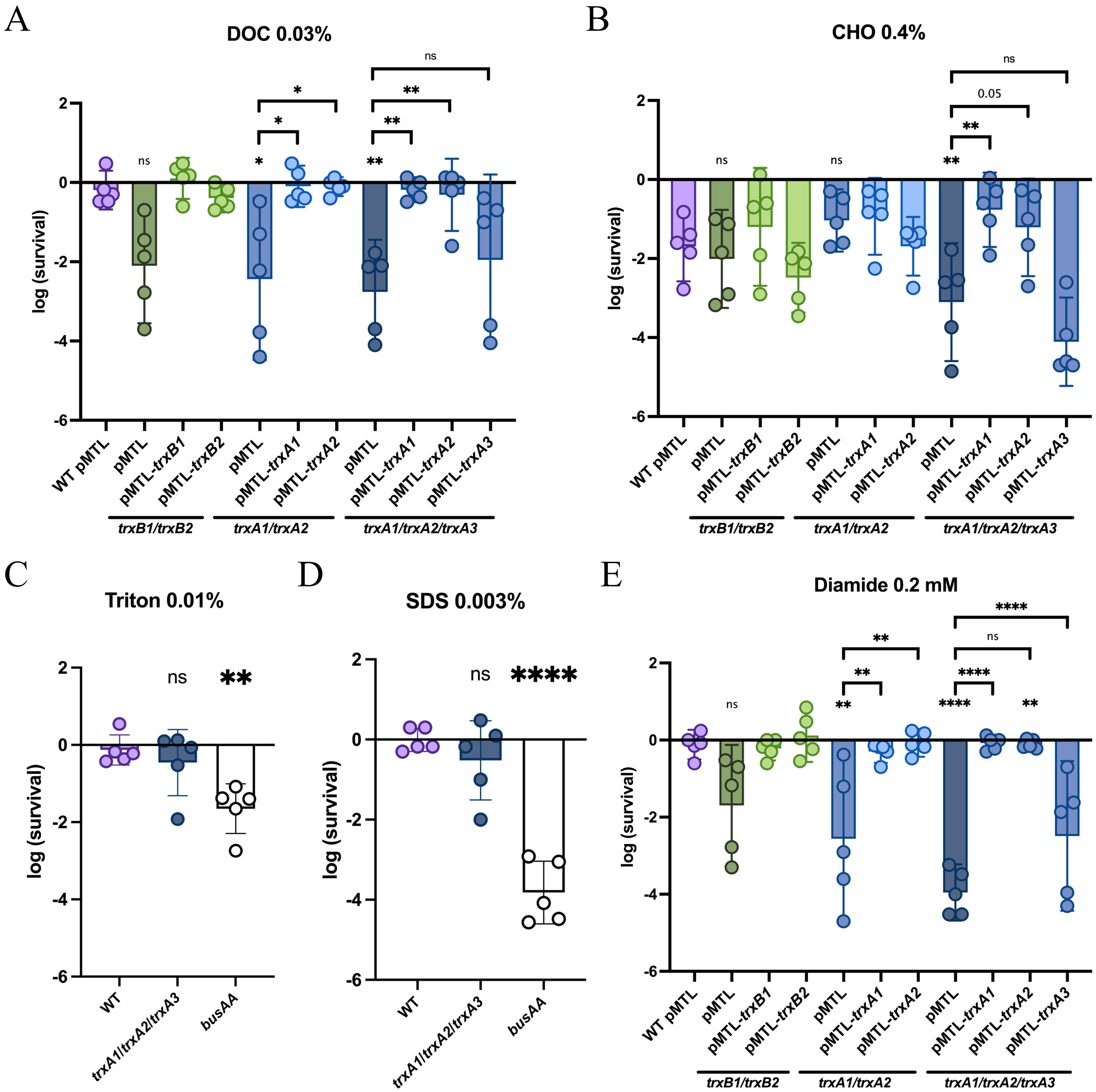

A
DOC 0.03%
B
CHO 0.4%
C
Triton 0.01%
D
SDS 0.003%
E
Diamide 0.2 mM
log (survival)
WT pMTL
pMTL
pMTL-trxB1
pMTL-trxB2
pMTL-trxA1
pMTL-trxA2
pMTL-trxA3
trxB1/trxB2
trxA1/trxA2
trxA1/trxA2/trxA3
WT
busAA
ns
0.05
